# High selectivity of frequency induced transcriptional responses

**DOI:** 10.1101/2022.10.06.511167

**Authors:** Alan Givré, Alejandro Colman-Lerner, Silvina Ponce Dawson

**Affiliations:** Departamento de Física, FCEN-UBA, and IFIBA, CONICET-UBA, (1428) Buenos Aires, Argentina; Department of Physiology, Molecular and Cellular Biology, School of Exact and Natural Sciences, University of Buenos Aires, Buenos Aires, Argentina; Institute of Physiology, Molecular Biology and Neurosciences, National Scientific and Technical Research Council (IFIBYNE-CONICET), Buenos Aires, Argentina

## Abstract

Cells continuously interact with their environment, detect its changes and generate responses accordingly. This requires interpreting the variations and, in many occasions, producing changes in gene expression. In this paper we use information theory and a simple transcription model to analyze the extent to which the resulting gene expression is able to identify and assess the intensity of extracellular stimuli when they are encoded in the amplitude, duration or frequency of a transcription factor’s nuclear concentration. We find that the maximal information transmission is, for the three codifications, ~ 1.5 – 1.8 bits, *i.e*., approximately 3 ranges of input strengths can be distinguished in all cases. The types of promoters that yield maximum transmission for the three modes are all similarly fast and have a high activation threshold. The three input modulation modes differ, however, in the sensitivity to changes in the parameters that characterize the promoters, with frequency modulation being the most sensitive and duration modulation, the least. This turns out to be key for signal identification. Namely, we show that, because of this sensitivity difference, it is possible to find promoter parameters that yield an information transmission within 90% of its maximum value for duration or amplitude modulation and less than 1 bit for frequency modulation. The reverse situation cannot be found within the framework of a single promoter transcription model. This means that pulses of transcription factors in the nucleus can selectively activate the promoter that is tuned to respond to frequency modulations while prolonged nuclear accumulation would activate several promoters at the same time. Thus, frequency modulation is better suited than the other encoding modes to allow the identification of external stimuli without requiring other mediators of the transduction.

## INTRODUCTION

Living organisms react to changes in their environment. The signaling systems that are used to “interpret” these changes and generate end responses are ubiquitous: they operate in both bacteria and eukaryotes and in processes as diverse as bacterial chemotaxis or the maturation of the immunse system, among many others. The malfunction of the information transmission systems involved in these processes is the cause of various pathologies. For this reason, understanding the way that living systems process and transmit information is fundamental from both a basic and applied point of view. The generation of responses to external stimuli usually involves changes in the intracellular concentration of some intermediaries which often induce changes in the nuclear concentration of transcription factors (TF) and, consequently, in gene expression. Sometimes, the intensity of the stimulus is encoded in the *amplitude* of this concentration, in others in the time it remains at a high level (*duration*) and in others in its *frequency* of variation [1–3]. Are there any advantages associated to a particular type of codification? When is one of these modes better suited than the other? These are two motivating questions of the present work.

In many cases, different external stimuli are encoded, transmitted and decoded by common signaling components [4–6]. In particular, the same TF can induce different end responses depending on the dynamics of its nuclear concentration. For example, p53 pulses induce the expression of DNA repair genes while a single sustained p53 pulse leads to the expression of senescence genes [7]. Something similar happens with the TFs, NF-kB, also in mammalian cells [8–10] and Crz1 in the yeast *S. cerevisiae* [11], and other genes. The TF, Msn2, which participates in the regulation of the multi-stress response in yeast [12, 13], is a good motivating example for the studies of the present paper. In response to glucose starvation, the nuclear translocation of Msn2 is oscillatory with frequency that is higher the lower the concentration of glucose. In response to osmotic stress, the time that Msn2 remains at a relatively high concentration in the nucleus increases and, in response to oxidative stress, it is the nuclear concentration of Msn2 the quantity that increases [2]. This suggests that promoters of genes induced by glucose deprivation (like DCS2 [14] and HXK1 [15]) are activated by *frequency modulation*, that promoters of genes induced during osmotic stress (*e.g*., SIP18 [16, 17] and ALD3 [18, 19]) are activated by *duration modulation* [20] and that those of genes induced by oxidative stress are activated by *amplitude modulation*. Following the works of O’Shea’s group [2, 20, 21] we may say that the type of codification identifies the type of stimulus. The question then arises as to how the promoters involved in these responses, which are regulated by the same TF, differ from one another so as to be induced by one or another type of modulation. This is another motivating question of the present study which, as the previous ones, can be addressed within the framework of information theory [22, 23]. This approach has been used to analyze cell signaling and infer properties of the underlying network of interactions, both from a theoretical point of view [24, 25] or using experimental data [21, 26, 27]. The information-based description involves defining what constitutes the input, the output and the channel connecting them. Using information theory it is possible to quantify the extent to which the output “distinguishes” the values that the input can take on. Given the distribution of input values, this is quantified by the *mutual information* between input and output, which is usually measured in bits, with *n* bits corresponding to distinguishing 2^*n*^ (sets of) input values. In the case of the cell, the ability to distinguish different inputs is key to generate adequate responses to each situation.

In this paper we analyze the mutual information between stimulus and response when the channel involves encoding the stimulus strength in either the amplitude, the frequency or the duration of a TF’s nuclear fraction. To this end we use the simple model introduced by Hansen and O’Shea to describe the activity of Msn2, defining as the input the amplitude, duration or frequency of the TF in the nucleus and, as output, the accumulated amount of mRNA produced, which correlates well with protein expression [20]. To account for stochastic fluctuations, which are not negligible in biological processes and can limit the information capacity of cell signaling pathways [26, 28, 29], we not only model the transcription step stochastically but also include noise in the amplitude of the TF concentration and in its interpulse frequency. Hansen and O’Shea [26] applied information theory to quantify the gene expression information transduced by Msn2 in yeast using experimental data obtained with high-throughput microfluidics. Instead, our approach is theoretical and seeks to determine the largest mutual information that can be achieved depending on the stimulus strength encoding. In addition, we ask whether there are disjoint regions of optimal parameter values for each encoding type that could allow the use of a single TF to elicit different end responses depending on the encoding. We found that, in most cases, the maximum possible mutual information is ~ 1.5 – 1.8bits, which is slightly larger than the values estimated from experiments in wild type yeast cells but similar to those obtained in mutants [21]. As discussed later, this value can be improved depending on the timing of the end response generation. We also found that the information transmitted, irrespective of the mode of encoding, is overall higher as the threshold for gene expression is higher or occurs on a faster timescale. Of the three modes, we found that frequency encoding is the most sensitive to changes in the kinetic parameters, while duration enconding is the least sensitive. This explains the experimental results of Hansen and O’Shea [21] and suggests that the cell can realize dynamic multiplexing by using a gene that transmits a large amount of information for the least sensitive modulation (duration) and a much smaller amount through the highly-sensitive channel (frequency).

## METHODS

### Model, inputs and outputs

We consider the simplified transcription model [20] depicted in Fig. 1. In this scheme, *TF* is the (nuclear) transcription factor which is assumed to undergo a known dynamics (piecewise constant in time). *P*_0_(*t*) and *P*_1_(*t*) represent the promoter in its inactive or active state, respectively (*P*_0_ + *P*_1_ = 1). The promoter is activated by *TF* in a cooperative fashion, as reflected in the term, 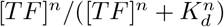, where *n* indicates the cooperativity and *K_d_* represents an effective dissociation constant of the binding/unbinding reaction which is assumed to occur on a faster timescale than the rest of the processes and, therefore, be in equilibrium. The *TF*-bound promoter then becomes active with rate, *k*_1_. The arrow from *P*_1_ to mRNA represents transcription which occurs at rate, 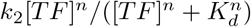, meaning that *TF* needs to be bound for transcription to take place. Given that 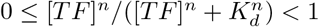, *k*_1_ and *k*_2_ are, respectively, the maximum rates at which promoter activation and transcription occur. The model includes mRNA degradation at rate, *d*_2_, but not the translation into the protein which is assumed to decay slowly enough so that the accumulated amount of mRNA produced can be used as the output of the process.

In the simulations, transcription is modeled stochastically with a master equation, while the rest of the steps are modeled deterministically, using Euler’s method to solve the ODEs. The *TF*’s nuclear concentration, [TF](t), is modeled by a single or a sequence of square pulses. The maximum TF amplitude considered is 100 (in dimensionless units) and the maximum pulse duration is 10 min. The total time of the simulations is 100min. The value, *P*_1_(*t*), derived from the numerical integration of the first two steps and [*TF*](*t*) are fed into the transcription Markov Process where the core of the randomness occurs. Finally, the mRNA is integrated in time to obtain the output. Given that *P_o_* + *P*_1_ = 1, the simulations imply solving the following equations:

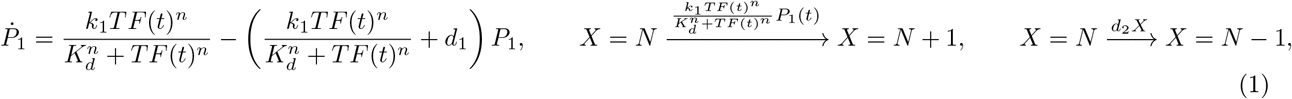

where *X*(*t*) is the (stochastic) number of mRNA molecules at time, *t*, and *N* = {0, 1, 2 ⋯}.

Three types of *TF* modulations are studied: duration (identified with the label “0”), amplitude (identified with the label “1”) and frequency (identified with the label “2”). In the cases of modulation by duration or amplitude, *TF*(*t*) is given by a single pulse of amplitude, 100, and variable duration or of variable amplitude and duration, 10*min*, respectively, that starts at *t* = 0. In the case of frequency modulation,*TF* pulses of 1*min* duration and amplitude, 100, that occur all throughout the simulation are considered with a Poisson process to determine their timing. In all cases, *TF*(*t*) is the norm of the sum of the deterministic pulse amplitude at time t and a normally distributed random variable of standard deviation, 10. The output is:

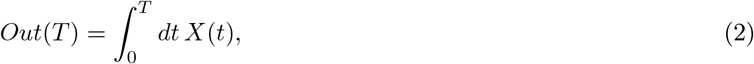

with, in most cases, one of three finite values, *T* = *T_a_, T_b_* and *T_c_*. These values are *T_a_* = 10*min, T_b_* = 50*min* and *T_c_* = 100*min* and *T_a_* = 1*min*, *T_b_* = 10*min* and *T_c_* = 100*min* for frequency and amplitude modulation, respectively. For the latter, *T_b_* corresponds to the time at which the *TF* pulse ends for all amplitudes. For duration modulation, we use *T_a_* = *t_end_, T_b_* = 10*min* and *T_c_* = 100*min*, with *t_end_* the duration of the input *TF* pulse. Notice that the *T_a_* time cut of this case is qualitatively different from the other 8 cuts, since it does not correspond to a fixed time, but instead it is defined by an event. This makes the promoters selected to be sensitive to that particular setting different from the others. For illustrative purposes we also compute, *Out*(*T*), and the corresponding mutual information, as continuous functions of time, *T*.

### Computation of Mutual Information

Discretizing the set of values that the input, *I*, and the output, *O*, can take on 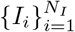 and 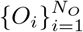, respectively), their mutual information can be written as:

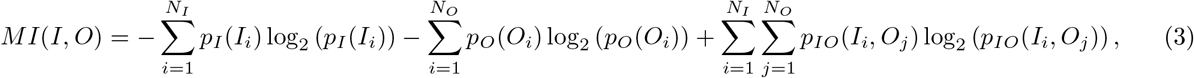

where *p_I,O_* is the joint probability distribution of *I* and *O* and *p_I_* and *p_O_* are the corresponding marginal distributions. All the simulations are done assuming a uniform distribution of the input values: over [1, 10]*min* for duration modulation; over [0,100] for amplitude modulation; over (0,0.1]/min for frequency modulation. The highest frequency considered gives, on average, a time integral of [*TF*](*t*), over the simulation time interval (100*min*), that is equal to the equivalent integrals for the cases of amplitude or duration modulation (100 × 10*min*). The uniform distribution implies that *p_I_*(*I_i_*) = 1/*N_I_* ∀ *i*, so that the first term in the r.h.s. of Eq. (3) is equal to log_2_(*N_I_*). All the simulations are done discretizing the corresponding input in *N_I_* = 200 values. Given the input parameters that are kept fixed depending on the modulation type (pulse amplitude and/or duration) and a set of model parameters (*k*_1_, *k*_2_, *n*, *K_d_*, *d*_1_ and *d*_2_), 15,000 simulations are run for each of the 200 input values that correspond to the modulation type probed (amplitude, duration or mean inter-pulse frequency). The results of these 3 10^6^ simulations are then used to compute the 3 types of output cuts described before (which only differ in the time interval over which the number of mRNA molecules is integrated, see Eq. (2)). The mutual information between each of the three input types and each of the three output cuts (9 combinations) is computed, for each set of kinetic parameter values, using the Jackknife method, which corrects for undersampling [30, 31]. For some studies we also compute *MI* as a function of time, in which case we call it *MI* (time). We compute *MI* (time) for each input type integrating *X* in Eq. (2) between 0 and time.

### Sampling of parameter space, optimization and comparisons

One of the aims of the present study is to determine the parameters of the model that maximize *MI* for each input modulation and output time cut. To guarantee a homogeneous sampling of the parameter space, the Latin Hypersquare Sampling (LHS) method was used [32], dividing the logarithmic range of values of each parameter (*k*_1_, *k*_2_, *K_d_*, *n, d*_1_) into equiprobable, non-overlapping intervals [33]. The intervals sampled for each parameter are shown in Table (I). In total, 17500 parameter sets were probed.

**TABLE I.** Sampled intervals for each of the parameters of the problem.

| Parameter | Range |
| --- | --- |
| $k_1$ (1/min) | (0.01,1) |
| $k_2$ (1/min) | (1,100) |
| $n$ | (1,10) |
| $K_d$ (a.u.) | (10,100) |
| $d_1$ (1/min) | (0.01,1) |
| $d_2$ (1/min) | 0.12 |

To obtain the results of the subsection “One transcription factor, two genes” we look for the set of parameters that gives the smallest value, *MI_m_*, for one type of modulation and output time cut, *i* (*i* = 0, 1, 2 as described before) restricting the search over the sets of parameters that give values, *MI*, that are within 90% of the maximum, *MI_M_*, for another type of modulation (*j* ≠ *i*) and the same time cut. We then define the *Minmax* region of parameters for the corresponding pair of input types, *ij*, as the sets that give values of *MI* that differ by less than 10% from *MI_M_* and *MI_m_* for the *i* and *j* input modulation types, respectively. We also say that the sets of parameters in the *Minmax* fulfill the *Minmax* condition.

## RESULTS

To analyze the mutual information (*MI*) between a stimulus and its induced transcriptional response, and how *MI* depends on whether stimulus strength is encoded in amplitude, frequency or duration of a transcription factor activation state, we performed simulations of a simple transcriptional model first presented by Hansen *et al* to describe gene induction by the TF Msn2 of the budding yeast *S.cerevisiae* [20]. As illustrated in Fig. 1, we assumed that increasing stimulus strengths could modulate nuclear TF (TF, the input of our model) in three possible ways: larger *TF* concentrations (amplitude modulation), longer periods of maximal *TF* concentration (duration), and higher frequency of *TF* pulses (frequency). The output (transcription) was obtained integrating the number of mRNAs produced over three different time intervals. Given the *TF*, each set of parameter values of the transcription model corresponds to a different promoter. In this Section we show the results obtained when *MI* was computed for each combination of input modulation and output time cut. In some instances, we also show the results of computing *MI* as a continuous function of the output time cut. To interpret the results, we analyze the time course of some key variables of the model. We study as well how *MI* varies with the parameters of the model and whether there are sets for which *MI* is large for an input modulation type and much smaller for another.

**FIG. 1.**
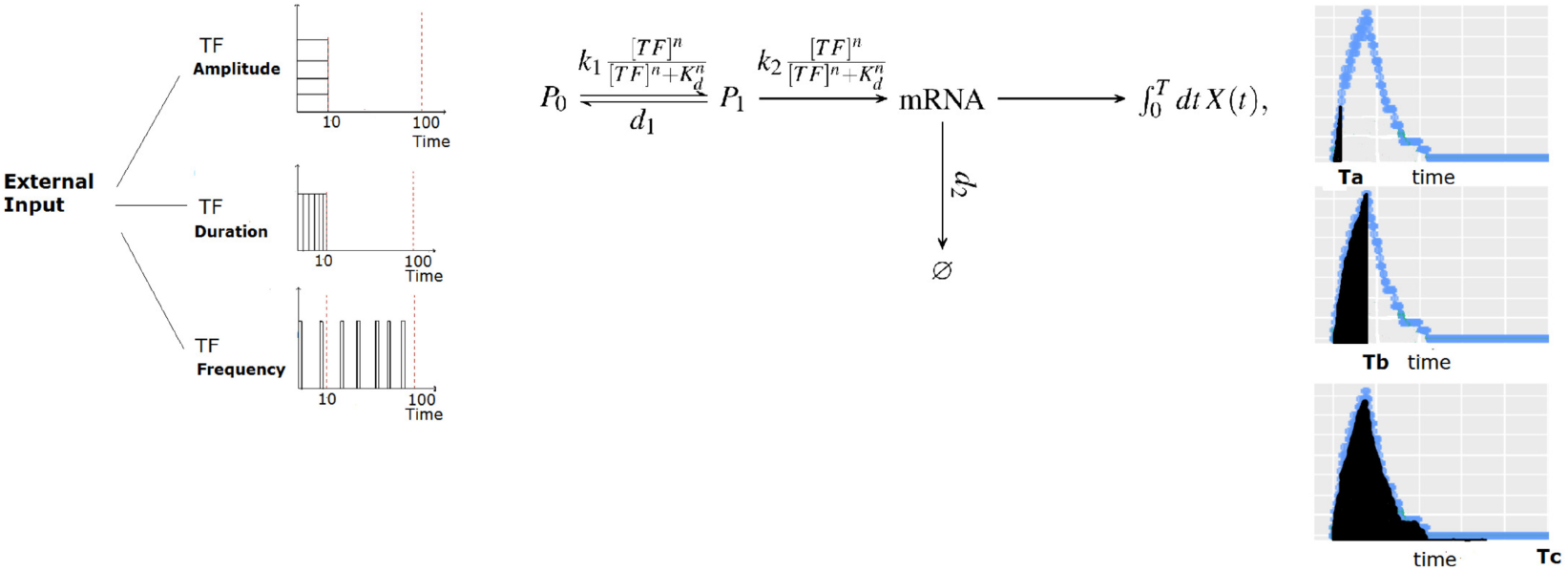
The model. An external stimulus is encoded in the (pulse) duration, (pulse) amplitude or (inter-pulse) frequency of the nuclear *TF* concentration, [*TF*]. The model does not describe this first step of the codification process, but starts directly with the nuclear *TF* time course. The mutual information, *MI*, between the input (pulse duration, pulse amplitude or mean inter-pulse frequency, depending on the type of stimulus codification or “TF modulation type”; 3 subfigures on the left) and the output (the time integral of the number of mRNA molecules produced, *X*) is computed choosing three possible time cuts for its calculation (3 subfigures on the right which correspond to an example with a single TF pulse of amplitude, 100, and 10 min duration). In between there is the transcription model [20] described in more detail in the main body of the paper

### Maximum mutual information and model parameters

In this subsection we show the results of the optimization run for each combination of input modulation type and of output time cut when five of the parameters of the model, *d*_1_, *k*_1_, *k*_2_, *K_d_* and *n*, are varied maintaining *d*_2_ constant at 0.12/*min* (see Sec.). The sets of parameters that maximize *MI* in each case and the range of values over which the search for the maximum was performed are shown in Table II. We see that *MI_M_* is between 1 and 2 bits for all combinations (with a value ~ log_2_(3), which is equivalent to distinguishing 3 values), with the exception of one special case that gives 2.6. Thus, in most cases, the system has, in principle, the possibility of behaving better than a binary (noisy) switch (1 bit). In the case of frequency modulation, we observe that *MI_M_* increases with the integration time from 1 to 2 bits, indicating that the more time allowed to the system to process the input, the more information it is able to extract. In constrast, for amplitude modulation, *MI_M_* behaves non-monotonically with the integration time, reaching almost 2 bits at the intermediate time that marks the end of the *TF* (single) pulse. The reduction in *MI_M_* at longer times in this case is likely due to loss of information during the noisy decay of the mRNA between the end of the stimulus and the end of the output integration time. Lastly, the case of duration modulation is significantly different from the other two cases in that when assayed at the shortest output integration time interval, the highest value, *MI_M_* = 2.61, is obtained. It must be pointed out, however, that in this case the output is integrated only while TF(*t*) is different from 0 (while the pulse is on). This is qualitatively different from the other cases for which we used a fixed time to integrate the output and is the main reason why we obtained a significantly larger value of *MI_M_* for *T_a_*.

**Table II.** Sets of parameters that maximize the mutual information, MI, for each combination of input modulation and Output integration time and the corresponding value, *MI_M_*, obtained in each case. The table shows the range over which each parameter value was varied when looking for this maximum. The parameter, *d_2_*, was always fixed at *d*_2_ = 0.12/*min*.

| Input Modulation | Output Cut | $k_1$ [1/min]<br>(0.01,1) | $k_2$[1/min]<br>(1,100) | $n$ [a.u.]<br>(1,10) | $K_d$ [a.u.]<br>(10,100) | $d_1$ [1/min]<br>(0.01,1) | $MI_M$ [bits] |
| --- | --- | --- | --- | --- | --- | --- | --- |
| Duration | $T_a$ | 0.32 | 91 | 1 | 10 | 0.056 | 2.61 |
| Duration | $T_b$ | 0.6 | 84 | 7.3 | 79 | 0.017 | 1.61 |
| Duration | $T_c$ | 0.13 | 63 | 8.3 | 94 | 0.016 | 1.52 |
| Amplitude | $T_a$ | 0.62 | 92 | 9.5 | 49 | 0.024 | 1.65 |
| Amplitude | $T_b$ | 0.93 | 31 | 3.3 | 96 | 0.13 | 1.91 |
| Amplitude | $T_c$ | 0.72 | 45 | 4.2 | 81 | 0.84 | 1.66 |
| Frequency | $T_a$ | 0.77 | 26 | 9.9 | 100 | 0.031 | 0.82 |
| Frequency | $T_b$ | 0.08 | 45 | 6.5 | 55 | 0.012 | 1.52 |
| Frequency | $T_c$ | 0.13 | 35 | 7.3 | 54 | 0.017 | 1.87 |

We observe in Table II that the parameters that maximize *MI* have various features in common for almost all combinations of input modulation type and output time cut. Namely, if we set aside the output cut, *T_a_*, for the duration modulation input type, which is qualitatively different from the others, we observe that *k*_2_ ≫ *k*_1_, *d*_1_, *d*_2_, implying that the timescale of transcription must be as fast as possible to guarantee a good information transmission. Second, *n* ≥ 3 in most cases and the dissociation constant of the TF binding/unbinding reaction is *K_d_* ≳ 50, of the same order of magnitude but smaller than the maximum *TF* concentration (100). The lower bound, 50, is also 5 times larger than the noise amplitude that is considered in the model (10). *MI* attains its maximum for *k*_1_ ≫ *d*_1_, a condition that guarantees that, at steady state, 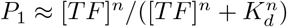. Thus, provided that *k*_1_ ≫ *d*_1_, the ability to distinguish different amplitudes that range between 0 and 100 will depend on how the function 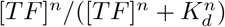 maps the [0,100] interval.

To analyze in more detail the effect of each parameter on *MI* we performed a sensitivity analysis varying each parameter while keeping the others fixed at the values that gave *MI_M_*. We show the results in Fig. 2. This figure confirms the initial conclusions that we drew from Table II. Namely, for all but one case (the duration modulation type for the output time cut, *T_a_*), *MI_M_* is attained when *d*_1_ is low and *k*_1_ and *k*_2_ are high, compared with the fixed timescale of the model, *d*_2_ = 0.12/*min*. If we look at the parameters related to the TF-promoter relationship, we observe that *K_d_* tends to be medium-high and the Hill Coefficient, *n*, tends to be high for the maximum *MI* to occur. On the other hand, we observe that the range over which the parameters can be varied without changing *MI* much is different ddepending on the parameter and the input modulation. In particular, we observe that amplitude modulation transmission is the most sensitive to separate changes in *K_d_* and *n*, that frequency modulation is most sensitive to changes in *d*_1_ while *MI* barely changes upon variations in *d*_1_ for duration modulation. In most cases, once the model parameters exceed a threshold, *MI* stays approximately constant and we observe, on average, the most extended plateaus for duration modulation.

**FIG. 2.**
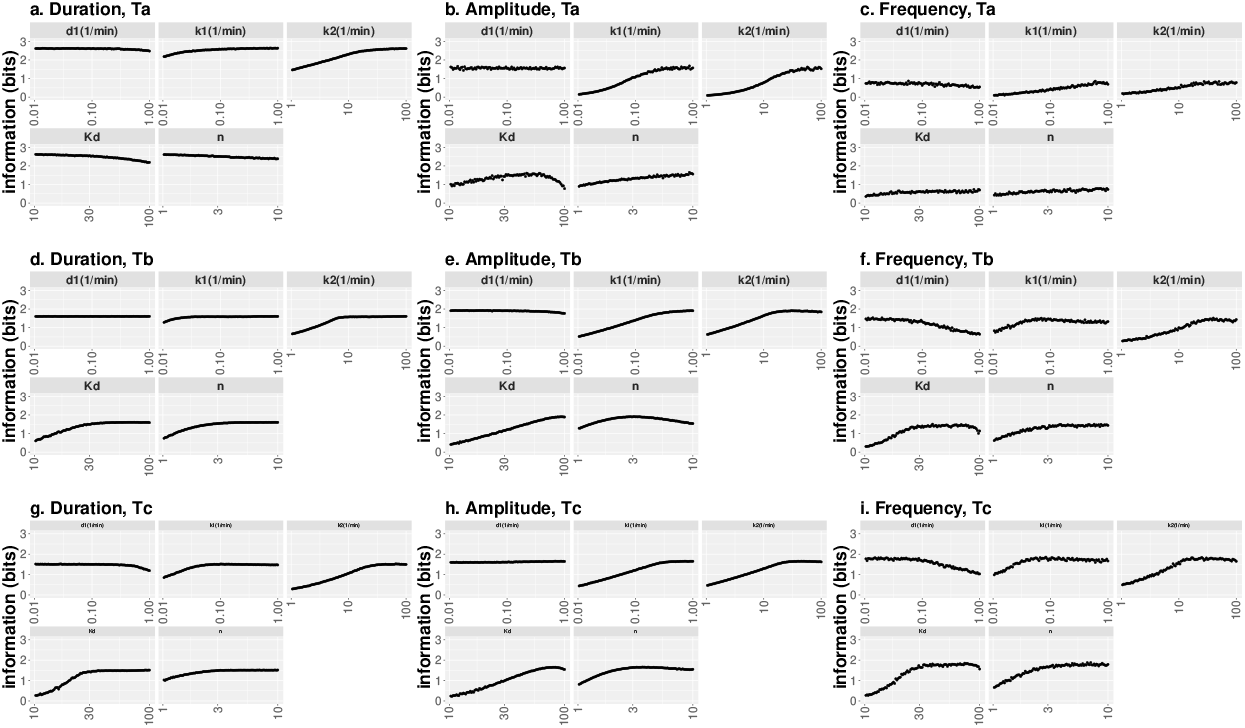
Behavior of mutual information, *MI*, around its maximum value, *MI_M_*, when one kinetic parameter is varied. We show *MI* as a function of the corresponding parameter for each type of input modulation and output time cut (**a**-**c**, **d**-**f** and **g**-**i** correspond, respectively, to the *T_a_, T_b_* and *T_c_* output time cuts; **a**-**d**-**g**, **b**-**e**-**h** and **c**-**f**-**i**correspond, respectively, to duration, amplitude and frequency input modulation type). Except for the modulation by duration and the *T_a_* time cut, the rest of the behaviors are very similar across modulations and time cuts. In particular, the highest information is obtained if *d*_1_ is low, *k*_1_, *k*_2_ and *n* are high and *K_d_* is in a middle-high range, but not too high. For the *T_a_* time cut and duration modulation type *K_d_* and *n* need to be low to maximize *MI*.

### Model dynamics and *MI* maximizing parameters

In order to understand why the maximum information transmission is obtained, in most cases, for parameters that satisfy *k*_2_ ≫ *k*_1_, *d*_2_, *d*_2_; *n* ≥ 3 and *K_d_* ≳ 50, we look at how the time course of some key variables of the model (*TF, P*_1_, mRNA and *Out*(*time*)) changes depending on whether the parameters are such that they give a relatively large or low value of MI for some combination of input type and output time cut.

We show in Figs. 3a-c the results obtained with simulations in which a single *TF* pulse of 10min duration was considered. When the parameters satisfy *k*_2_ ≫ *k*_1_, *d*_1_, *d*_2_, *n* ≥ 3 and *K_d_* ≳ 50 we observe that the three input strengths translate into three distinguishable outputs (Fig. 3a). In the other examples, either promoter activation dynamics is too slow and it never gets activated (Fig. 3b, where *k*_1_ ≪ *d*_1_) or the promoter is activated maximally for all inputs (Fig. 3c, where the threshold determined by *K_d_* and n is too low and the timescale is not slow). In the last case, the three input strengths translate into almost indistinguishable outputs. We show in Fig. 3 d the time course of *MI* computed for amplitude modulation and the *T_c_* output time cut using the kinetic parameters of the example in a. We observe that *MI* reaches a maximum approximately at *t* = 10*min, i.e*., when the TF pulses end, and then decays towards a slightly lower value.

**FIG. 3.**
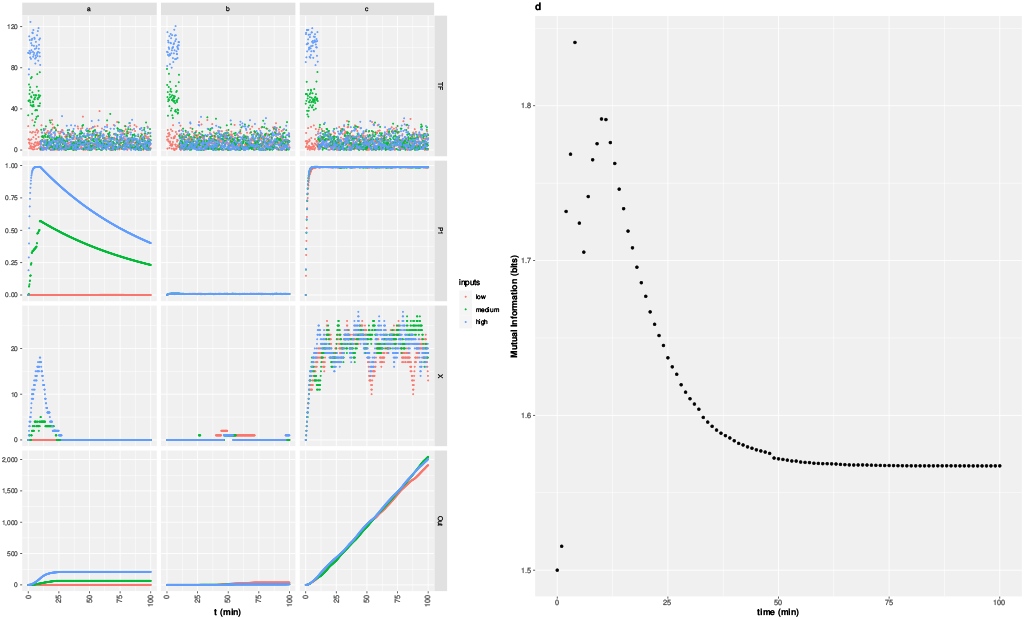
Time course of [*TF*], *P*_1_, *X* and *Out*(*time*) for 3 sets of parameter values and 3 choices of *TF*(*t*) (a-c) and time course of *MI* for amplitude modulation, the *T_c_* time cut and the kinetic parameters of the example in a. (a-c) In all the simulations, a single TF pulse of 10*min* duration was considered. The amplitudes were 0 (red), 10 (green) and 100 (blue) with a noise of amplitude, 10, superimposed in each case. The parameters used in the simulations of the first column were *k*1 = 1/*min*, *k*2 = 100/*min*, *d*1 = 0.01/*min*, *Kd* = 70, n = 10 for which the mutual information, computed for the amplitude modulation input type and the output time cut, *T_c_*, is *MI* = 1.56 (*i.e*. 94% of *MI_M_*). Those of the second column were *k*1 = 0.01/*min*, *k*2 = 1/*min*, *d*_1_ = 1/*min, K*_d_ = 10, n =1 for which *MI* = 0.03, and those of the third: *k*1 = 1/*min*, *k*2 = 100/*min*, *d*1 = 0.01/min, *Kd* = 10, *n* = 1 for which *MI* = 0.02. (d) *MI* as a function of time computed for amplitude modulated inputs and the *T_c_* time cut using the kinetic parameters of the example in a.

### Pairs of parameters and mutual information

So far, we have analyzed how *MI* varies when a single parameter is varied. In this Section we analyze if the parameters are interdependent in determining *MI_M_*. To this end, we varied the values of all possible pairs of parameters maintaning the others fixed at the values that gave *MI_M_* and calculated *MI* (Fig. 4). From the analysis of the results, we determined how the pairs had to be varied to keep *MI* constant. We found that, for all input and output types, *d*_1_ tended to be positively correlated with *k*_1_ and *k*_2_. That is, changing *d*_1_ in a given direction modifies *MI* in a way that is compensated by modifying *k*_1_ or *k*_2_ in the same direction. We then found that *k*_1_ and *k*_2_ were negatively correlated with one another, i.e., if *k*_1_ is increased, *k*_2_ has to be increased to keep *MI* fixed, and viceversa. Similarly, the parameters related to the *TF*-promoter relationship, *K_d_* and n, are negatively correlated in most cases, with the exception of the *T_a_* output time cut for the modulation by duration input type, for which *n* and *K_d_* need to be low for *MI* to attain its maximum value.

**FIG. 4.**
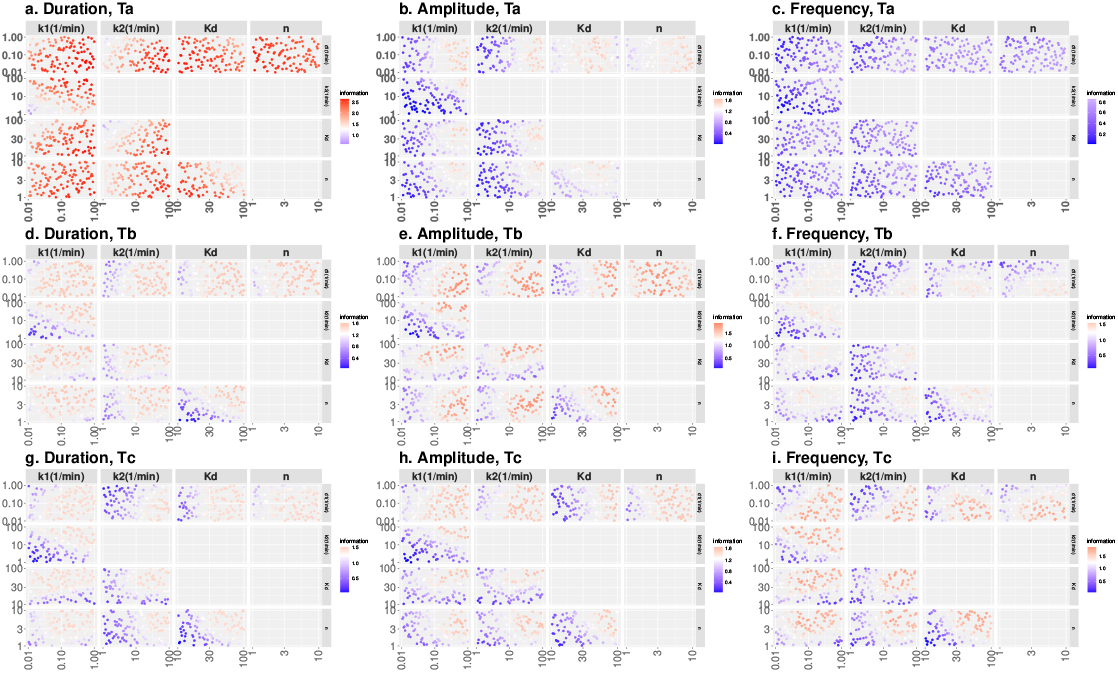
Mutual information, MI, as a function of two parameters while the others are left fixed at the values that maximize *MI* for the particular input and output types analyzed in each case (**a**-**c**, **d**-**f** and **g**-**i** correspond, respectively, to the *T_a_, T_b_* and *T_c_* output time cuts; **a**-**d**-**g**, **b**-**e**-**h**and **c**-**f**-**i** correspond, respectively, to duration, amplitude and frequency input modulation type). Common behaviors are observed throughout most modulations and time cuts.

One of the aims of the present work is to determine whether there could be two promoters that are modulated by the same *TF*, but each discriminates among stimulus strengths that are encoded in a different property of the nuclear *TF* concentration while, at the same time, is not good at discriminating the strengths encoded in the property at which the other one is good. For example, one promoter could be good at discriminating increased TF amplitudes but be “blind” to increases in frequency while the reverse situation be valid for a second promoter. To explore this possibility, we searched for the sets of kinetic parameters that gave *MI* within 90% of the maximum, *MI_M_*, for a certain input modulation type and, at the same time, gave a relatively small value for another input type (and the same output time cut), *i.e*., the sets that fulfill the *Minmax* condition (see Methods).

Fig. 5 shows the projection of the six *Minmax* regions, for the *T_c_* time cut, on each of the five parameter space axes of the transcription model. Although it should be analyzed with care because the one-dimensional projection of a 5-dimensional volume does not reflect its folds, Fig. 5 illustrates how much each parameter can be varied while satisfying the *Minmax* condition. In particular, we can observe that the sensitivity of *MI* to parameter variations is very different depending on the parameter and the pair of input modulation types. For example, *k*_1_ and *d*_1_ can be varied by over an order of magnitude and, yet, *MI* differs by less than 10% from *MI_M_* for the duration modulation type and from the conditional minimum, *MI_m_*, for the amplitude modulation type, and *viceversa*. The parameter, *d*_1_, on the other hand, can barely be varied if we try to keep *MI* within 90% of *MI_M_* for the amplitude or the duration modulation type and differ by less than 10% from *MI_m_* for the frequency modulation type. Overall, we see that keeping *MI* as small as possible for frequency modulation in each *i*2 *Minmax* region requires a very fine tuning of the time-related parameter values (*k*_1_, *k*_2_ and *d*_3_) and, in the case of the 12 region, of the dissociation constant, *K_d_*, as well. As we show in what follows, this fine tuning allows the finding of promoters (characterized by sets of parameter values) that are “blind” to frequency modulated inputs but are good at transmitting information in other modulation types. The reverse situation is impossible or very hard to find for the duration or the amplitude modulation, respectively.

**FIG. 5.**
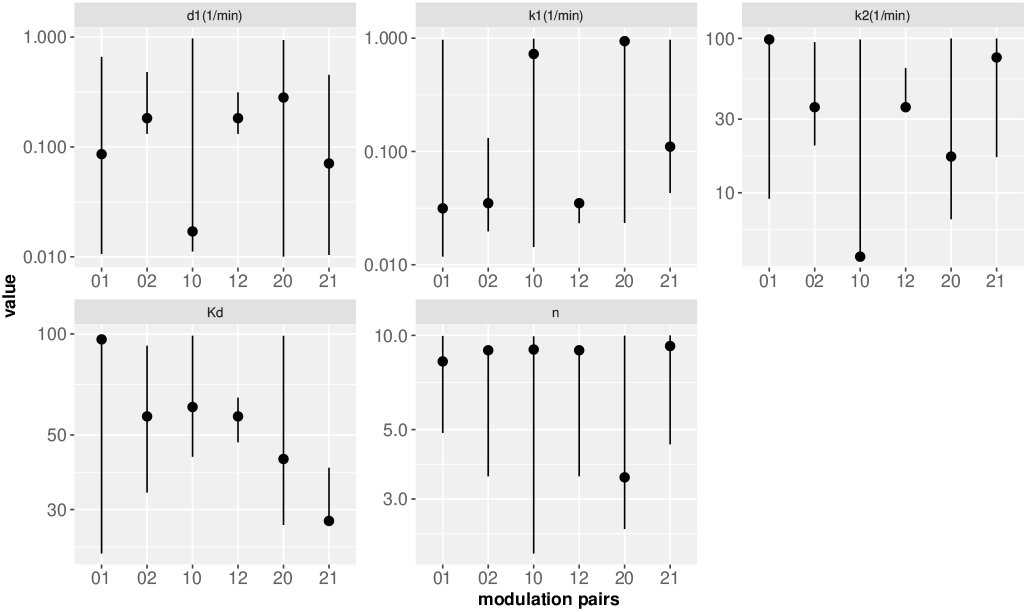
Projection on each parameter axis of the *Minmax* regions obtained for each pair of modulation input types and the *T_c_* output time cut. The pairs of digits in the horizontal axes identify the modes for which the information transmission has been maximized (first digit) and subsequently minimized (second digit) to determine the *Minmax* regions, with 0: duration, 1: amplitude and 2: frequency. The symbols correspond to the values that minimize *MI* for the second digit modulation given that MI stays within 90% of *MI_M_* for the first digit modulation.

We now analyze the difference in information transmission that is achieved when choosing some of the model parameters determined with the *Minmax* conditioning. This is illustrated in Fig. 6a where we show the values of *MI* obtained when the information is computed for each transmission mode for one set of parameters in the *Minmax* region that corresponds to the pair of compared transmission modes. The parameters chosen are those that minimize the information through one of the modulations while still being within 90%of the other’s maximum (plotted with circles in Fig. 5). We observe that *MI*, for the duration modulation type, has approximately the same value (~ 1.3 –1.4) for all combinations, including those in which *MI* by duration was minimized. Given that each set of parameter values corresponds to a different promoter, this implies that durations would always be equally discriminated regardless of the promoter. In contrast, frequency modulation is the most sensitive mode with *MI* variations of almost one bit depending on the parameters that characterize the promoter (MI ~ 0.8 for the sets depicted in red or blue and *MI* ~ 1.7 for those depicted in orange or cyan). The situation for the amplitude modulation type is intermediate with variations of about 0.5 bits (*MI* ~ 1.1 for the parameters depicted in brown or cyan while *MI* ~ 1.5 for those depicted in green or blue). These results indicate that it should be possible to find promoters that, being regulated by one TF, are good at decoding duration or amplitude encoded inputs and, at the same time, be “blind” to frequency modulated ones, but that the separation of behaviors would not be as clear in the opposite situation, especially in the case of the combination frequency-duration. We now analyze how the discriminating ability is reflected in the amount of mRNA that is produced using some examples (Fig. 6).

**FIG. 6.**
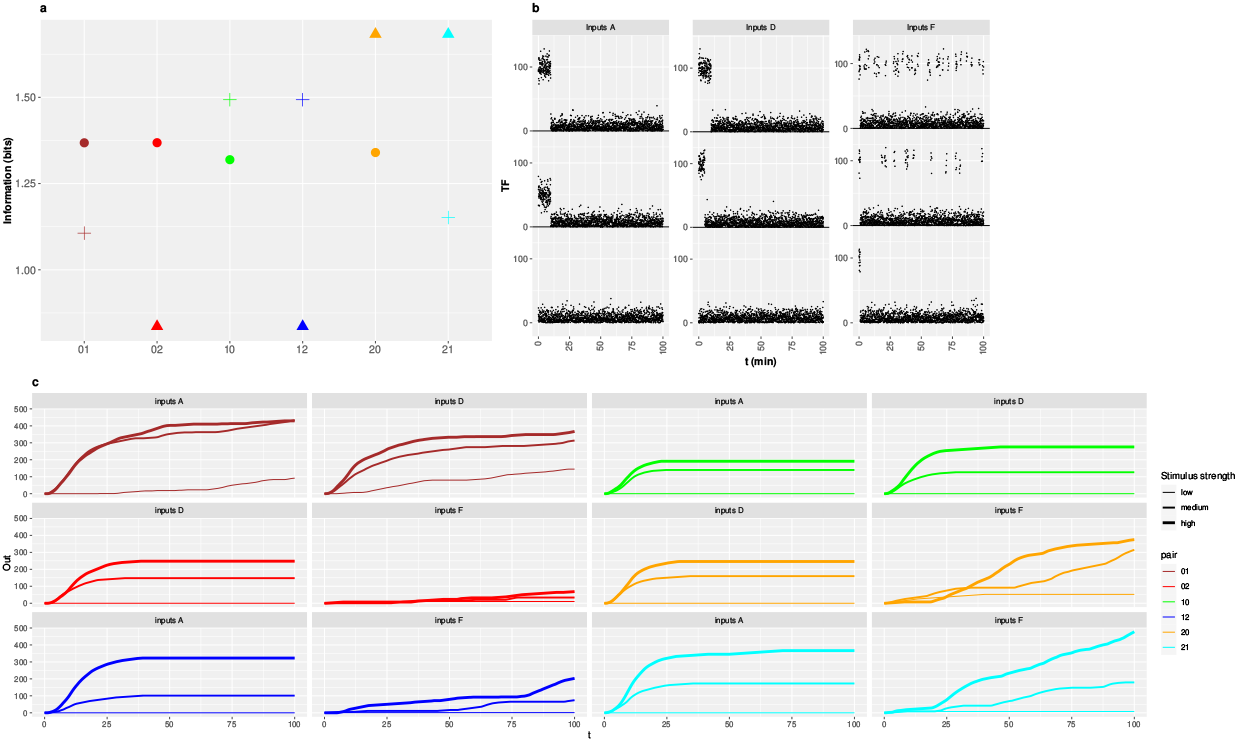
Mutual information and dynamical behaviors obtained using the set in each *Minmax* region that minmizes MI for the “second” modulation type. **a**: MI computed using the selected set of parameters in each *Minmax* region for the input modulation modes that are maximized and conditionally minimized to determine the region (circles: duration, crosses: amplitude, triangles: frequency) and the *T_c_* output time cut. The labels on the horizontal axis identify the *Minmax* regions with the digits order as in Fig. 5. Each parameter set is identified by a different color. The parameter values in each set are displayed with symbols in Fig 5.**b-c**: Accumulated mRNA (*Out*(*time*)) as a function of time (**c**) obtained from simulations of the model using the *TF* time courses displayed in **b** (curves in **c** are plotted with increasing thickness with stimulus strength) and the same parameter sets (identified by the same color) as in **a**.

We show in Fig. 6c the mRNA that is accumulated as a function of time for various examples that were obtained using the parameter sets of Fig. 6a. We probed the dynamics of the model using these sets of parameters (each one of which can be associated with a different promoter that responds to the same *TF*) and sets of inputs, *TF*(*t*)-A, B, C, characterized, respectively, by decreasing duration, amplitude and frequency of the TF pulse or pulses, as shown in Fig. 6b, related to the transmission modes that had been maximized and conditionally minimized to determine the parameter sets.

Consistent with the previous discussion (Fig. 6a), it is the frequency encoding (Inputs F) the one that produces the largest differences depending on the parameter set. Not only the amount of mRNA produced for Inputs F is much larger for the two sets in which transmission by frequency modulation was maximized (20 and 21 *Minmax* regions) than in those in which it was conditionally minimized (02 and 12), but also the three TF time courses within Inputs F are more easily distinguishable in the former than in the last two cases. In the case of duration encoding (Inputs D), the amounts of accumulated mRNA are slightly larger for the set that belongs to the 01 *Minmax* region than for the one that belongs to the 10 one and slightly larger for the latter in comparison with the ones in the 02 or the 20 *Minmax* regions. The ability to distinguish the 3 durations does not seem to vary much with the parameter set. In the case of amplitude encoding (Inputs A), the amount of accumulated mRNA is largest for the set which was determined by conditionally minimizing the information transmission for amplitude modulation. For this type of inputs, the failure to transmit enough information for the parameter set in the 01 *Minmax* region compared with the set in the 10 one (see brown *vs* green cross in Fig. 6a) seems to be related to producing similar amounts of mRNA regardless of the *TF* amplitude in the former (see brown *vs* green curves for Inputs A). Something similar occurs when the behavior obtained for Inputs A and the sets in the 12 and the 21 *Minmax* regions are compared (blue and cyan curves of Inputs B, respectively).

## DISCUSSION AND CONCLUSIONS

In this paper we have used information theory to analyze the capabilities to transmit information via transcription when the extracellular stimuli are encoded in the amplitude, duration or frequency of the TF’s nuclear concentration. We used the simple model of Fig. 1-introduced by Hansen and O’Shea [20] to describe the activity of Msn2 in yeast-in which each promoter that is regulated by the same TF corresponds to a different set of parameter values of the transcription step. We computed the mutual information (MI) between the amplitude, the frequency or the duration of the TF’s nuclear concentration (the input) and the accumulated amount of mRNA produced (the output) to determine both the maximum *MI* for each encoding and the range of parameter values (*i.e*., the type of promoters) that gives the maximum in each case. Our studies showed that, for any combination of physiologically feasible parameter values and any modulation input type, the maximum *MI* is between one and two bits (Table II), with the only exception of the *T_a_* output time cut and duration modulation input, which is qualitatively different from the rest and for which it could be ~ 2.6bits. The maximum values obtained for the other cases were slightly larger than those encountered in experiments on Msn2-regulated gene expression in wild type (WT) yeast cells [21] which showed that it operated as a noisy switch (transmitting ~ 1.1 – 1.3bits) for two genes, *HXK1* and of *SIP18*, when they were modulated, respectively, by frequency and amplitude. When the authors mutated the promoters with the aim of improving information transmission, *MI* for amplitude modulation increased to ~ 1.5bits [21], very similar to the maximum values that we found with our parameter exploration. Looking at the time dependence of *MI* for the parameters that gave a relatively good information transmission we observed that, for duration and amplitude modulation, *MI*(*t*) was maximum approximately when the longest pulse ended and then decayed to a slightly smaller asymptotic value. For these modulation types, the timing of the maximum and the timescale of the subsequent decay of *MI* are related to the moment at which mRNA production ends and to the mRNA degradation rate, d_2_, respectively, as illustrated in Fig. 3 d which corresponds to a case in which all pulses end at *t* =10*min* and 1/*d*_2_~ 8min. The non-monotonic behavior of *MI* with time is probably the reason that underlies the much larger *MI* value obtained for the duration modulation input and the *T_a_* output time cut which corresponds to the end of mRNA production in all cases and, thus, does not include the subsequent mRNA degradation steps (included in all other output time cuts). It is also consistent with the observation that *MI* increases steadily with time for frequency modulation, given that, in this case, the TF pulses occurred thoughout the simulation time. Perhaps, cells could take advantage of the non-monotonic time course of *MI via* a pre-equilibrium sensing mechanism [34], something that we have not explored in the present paper.

The model we have used is characterized by four parameters directly associated to timescales (*k*_1_, *k*_2_, *d*_1_ and *d*_2_, which we kept fixed at *d*_2_ = 0.12/*min*) and by another two (*K_d_* and *n*) related to the *TF*-promoter relationship. We found (Table II) that those that maximized *MI* for all combinations of input modulations and output time cuts corresponded to a fast transcription timescale (*k*_2_ ≫ *k*_1_, *d*_1_, *d*_2_ with *k*_1_ ≫ *d*_1_ in most cases). This is so because a high transcription rate (at constant mRNA degradation rate, *d*_2_) creates ample and input-sensitive mRNA fluctuations (Fig. 3). This result agrees with those of Hansen and O’Shea [20] in that a fast timescale generates better responses. Regarding the *TF*-promoter relationship, again we found a distinguishing result for the *T_a_* time cut and duration modulation input. In particular, we found that a larger dynamic range (lower *K_d_* and *n*) was favored in this case compared to the others. Restricting the comparison to the *T_c_* output time cut, we obtained the sharpest *TF*-promoter relationship (n ~ 7 –8) for the duration and the frequency input modulations and values, *K_d_*, that were several times larger than the noise amplitude, 10, and varied between 50% (for frequency modulation) and 94% (for duration modulation) of the maximum *TF* concentration, 100. Some of these findings agree with and others seem to differ from the experimental results of Hansen and O’Shea [20]. Namely, these authors classified Msn2-activated promoters as high (H) or low (L) treshold (requiring a high or a low Msn2 concentration for induction, respectively) and as fast (F) or slow (S) (induced quickly or requiring a longer time with Msn2 bound to induce transcription, respectively). Their experiments showed that high treshold traits were accompanied by slow timescales (*HS promoters*) and that these genes were those modulated by duration (i.e., osmotic stress). They found, in turn, that low treshold traits were coupled to fast timescales (*LF promoters*) and that these genes were those modulated by frequency (i.e., induced during glucose starvation). While (for the *T_c_* output time cut) we found, in agreement with these results, the maximum *MI* at a larger *K_d_* (*i.e*. a larger threshold) for the duration modulation than for the frequency one (albeit within the same order of magnitude), for the transcription timescale (determined by the ratio,) we found, opposite to the observations, a value twice as large for duration modulation compared to frequency modulation.

We can explain the differences between the optimal values derived from our study and those calculated with experimental data obtained from WT cells by Hansen and O’Shea [21] in terms of the different sensitivity that *MI* displays to parameter variations for the various input modulation types. In general, once the model parameters surpass a threshold, *MI* stays approximately constant (Fig. 2). On the other hand, in most cases the threshold for one parameter depends on the other parameter values, *e.g*., the *k*_2_ threshold decreases for increasing *k*_1_ (Fig. 4). Thus, the “optimal” parameters are not that meaningful *per se*, in the sense that they could be varied (especially, in pairs) without much change in *MI*. Our study showed that the range over which the parameters could be varied without changing *MI* much was different for the different parameters and input modulation types. For example, for the *Tc* time cut we obtained that amplitude modulation transmission was most sensitive to changes in *Kd* and n, that *MI* for frequency modulation varied by over half a bit if *d*_1_ or *k*_1_ did not stay within the same order of magnitude as the values that gave maximum transmission for this mode and that duration modulation was the least sensitive to simultaneous variations in *d*_1_ and *k*_1_ (Figs.2-4). This different sensitivity was clearly reflected in the results of the search of parameters that could give good transmission for one input modulation and a relatively low one for another (Figs. 5–6a). Namely, we determined that frequency modulation was so sensitive to parameter changes that promoters could be found which yielded a relatively large information transmission for duration or amplitude modulation and a much lower one for frequency modulated inputs: one which gave *MI* ~ 1.4bits for duration modulation and *MI* ~ 0.8bits for frequency modulation (Fig. 6a, red symbols) and another which gave *MI* ~ 1.5bits for amplitude modulation and *MI* ~ 0.8bits for frequency modulation (Fig. 6a, blue symbols). These values are very similar to those obtained by Hansen and O’Shea [21] using a mutated version of the promoter, SIP18, that is induced upon prolonged nuclear accumulation of Msn2 (see their Fig. 2). Expanding our search beyond the limits of the already defined *Minmax* regions, we found a set of parameters (i.e., a promoter) which gave *MI* ~1.1bits for duration, *MI* ~1.2bits for amplitude and *MI* ~0.7bits for frequency modulation. The last two values are closer to those obtained for SIP18 in WT cells. We think that the information transmission for frequency modulation could have been reduced even further had we used a more sophisticated (and realistic) model than the very simple one of Fig. 1. Hansen and O’Shea [21] asked why the cell should not “fine-tune the expression level of stress genes to the stress intensity”. Our studies seem to indicate that, at least in the case of SIP18, the parameters might have been tuned to make it as blind as it could be for frequency modulation without losing the capability of distinguishing ON from OFF without error in the case of prolonged nuclear accumulation of Msn2. In any case, according to our studies, even under optimal promoter parameters the information transmission could only be slightly better than 1bit, i.e., each single promoter cannot act as a rheostat. A similar process might underlie dynamic multiplexing in gene expression elicited by p53 [7], something that could be investigated introducing mutations that could change the activation threshold or characteristic timescales of the promoters involved. Our studies also show that the lower sensitivity to parameter variations of the duration and amplitude modulated inputs might have prevented this total blindness to be attained for the promoter, HXK1, which is physiologically induced by Msn2 pulses.

## ACKNOWLEDGEMENTS

This work was partially supported by UBA (UBACyT 20020170100482BA) and ANPCyT (2015-3824, 2018-02026).

